# GLYCAM Bacterial Carbohydrate Builder: a web-tool for modelling 3D structures of bacterial glycans

**DOI:** 10.64898/2026.01.26.700271

**Authors:** Ferran Nieto-Fabregat, Oliver C Grant, Xiaocong Wang, Daniel Wentworth, Robert J Woods, Roberta Marchetti

## Abstract

Here we present the GLYCAM Bacterial Carbohydrate Builder (https://glycam.org/cb), an enhanced version of the GLYCAM-Web Carbohydrate Builder^1^ structure modeller that integrates support for modelling bacterial glycans, enabling the straightforward generation of three-dimensional structural models. The tool integrates bacterial monosaccharide parametrisations into a curated, user-friendly web-based resource. It provides an intuitive interface for the generation of carbohydrate sequences and generates 3D structural models in PDB file format, as well as the input files required for performing molecular dynamics simulations with the AMBER software package. The current implementation includes a library of 18 bacterial monosaccharides, which can be used in combination with the already parametrised eukaryotic sugars to construct complex bacterial glycans. Common derivatives, including acetylation, methylation, and sulfation are also supported.

By validating and integrating bacterial sugar parameters into the GLYCAM-Web Carbohydrate Builder, this work reduces the technical barriers associated with bacterial glycan modelling and facilitates computational studies of complex bacterial glycoconjugates.

## INTRODUCTION

In eukaryotic glycans, the commonly found monosaccharides include relatively few types; hexoses, aminohexoses, deoxyhexoses, pentoses, and the well-known family of sialic acids. In contrast, bacteria possess an incredibly high number of monosaccharidic constituents.^2, 3^ These include, among the others, uncommon aminosugars, hexuronic acids, heptoses, octulosonic and nonulosonic acids, branched monosaccharides, and monosaccharides with noncarbohydrate substituents, resulting in peculiar and unique structures (Figure 1) which play crucial roles not only for the bacterial survival but also for their interaction with the external environment, affecting *inter alia* the human health and several therapeutic interventions. These bacterial glycoconjugates include: the peptidoglycan (PG),^2, 4^ the lipopolysaccharides (LPSs),^5, 6^ capsular- and exo-polysaccharides (CPSs and EPSs).^7^

**Figure 1.**
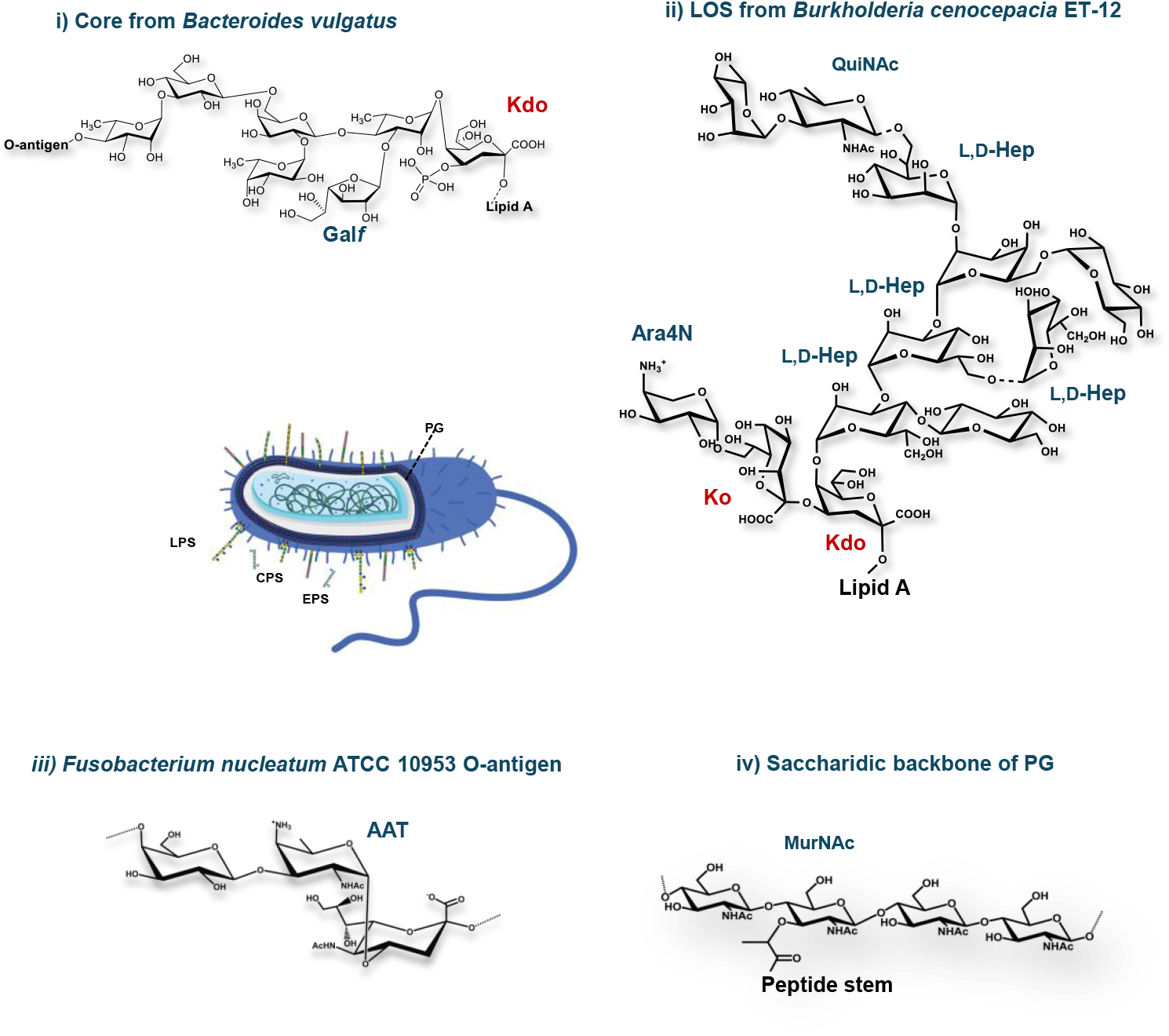
Schematic representation of the main glycoconjugates exposed on bacterial cell surfaces. The chemical structure of selected bacterial glycoconjugates, i) core OS from *Bacteroides vulgatus*,^8-10^ ii) LOS from *Burkholderia cenocepacia* ET-12,^11^ iii) O-antigen from *Fusobacterium nucleatum* ATCC 10953.^12^ iv) general disaccharidic backbone of PG, including unique monosaccharides, such as MurNAc, Kdo, Hep, QuiNAc, Ara4N, Gal*f*, and AAT, is also reported.

PG, one of the major structural components of bacterial cell walls, is a large polymer composed of alternating units of N-acetylmuramic acid (MurNAc) and N-acetylglucosamine (GlcNAc) β-1,4-linked, cross-linked by short peptide stems.^2, 4^

In Gram-negative bacteria, this scaffold is complemented by the presence in the outer membrane of LPS, which is composed of three main domains: the lipid A, the core oligosaccharide (OS) and the O-antigen polysaccharide (OPS).^5, 6^ The glycolipidic portion of the LPS, the lipid A, is generally composed of a bisphosphorylated glucosamine disaccharide backbone variously acylated and phosphorylated. In some bacteria, this conserved structure can be further modified, i.e. with the incorporation of a residue of 4-amino-4-*deoxy*-L-arabinose (L-Ara4N).^13^ The lipid A is linked to the core oligosaccharide by a peculiar sugar, namely Kdo (3-deoxy-D-*manno*-oct-2-ulopyranosonic acid), which is a marker of Gram-negative bacteria. This peculiar sugar moiety usually carries a negatively charged unit, such as another Kdo residue, or a D-*glycero*-α-D-*talo-*oct-2-ulopyranosonic acid (Ko) moiety, at its position 4. The core OS usually contains 6-10 monosaccharides including heptoses and hexose residues in addition to Kdo/Ko. In some microorganisms, as numerous bacteria belonging to the human gut, particular forms of sugars, as the galactofuranose, are also present.^8-11^ The polymeric O-antigen is characterized by a remarkable structural and chemical diversity, often entailing rare sugars.^13, 14^

Beyond the cell envelope, CPSs and EPSs are high molecular weight polymers, associated to the bacterial surface or secreted into the extracellular milieu, respectively, which protect bacteria from environmental stress and provide peculiar properties to the bacterial cells depending on their chemical structures.^7^

Interestingly, several feared pathogens expose on their surfaces glycans that mimic eukaryotic SAMP (*self*-associated molecular patterns) molecules, thus featuring sialic acid-like structures, such as legionaminic and pseudaminic acids, to exploit inhibitor host receptors and escape immune surveillance.^15^

Given the extraordinary diversity and chemical complexity of a high number of monosaccharides composing bacterial structures, modelling bacterial glycans is a challenging process. In practice, this diversity has often required the parametrisation of entire bacterial glycan ligands, or of specific monosaccharides and motifs within them, as they are encountered in individual biological systems. This process is time-consuming and has traditionally been performed on a case-by-case basis within single studies, frequently tailored to a specific organism or glycoconjugate. As a result, validated bacterial monosaccharide parameters are often dispersed across the literature, limiting their reuse and increasing the technical barrier for non-expert users.

Motivated by the growing accumulation of individually parametrised and experimentally supported bacterial sugars, we sought to consolidate these parameters into a curated and accessible resource. This effort led to the integration of a library of bacterial monosaccharides into the GLYCAM-Web Carbohydrate Builder, enabling their straightforward use within an established modelling framework. The resulting GLYCAM Bacterial Carbohydrate Builder (Figure 2) provides a user-friendly interface that allows both expert and non-expert users to build complex and biologically relevant glycan sequences through a point-and-click mechanism.

**Figure 2.**
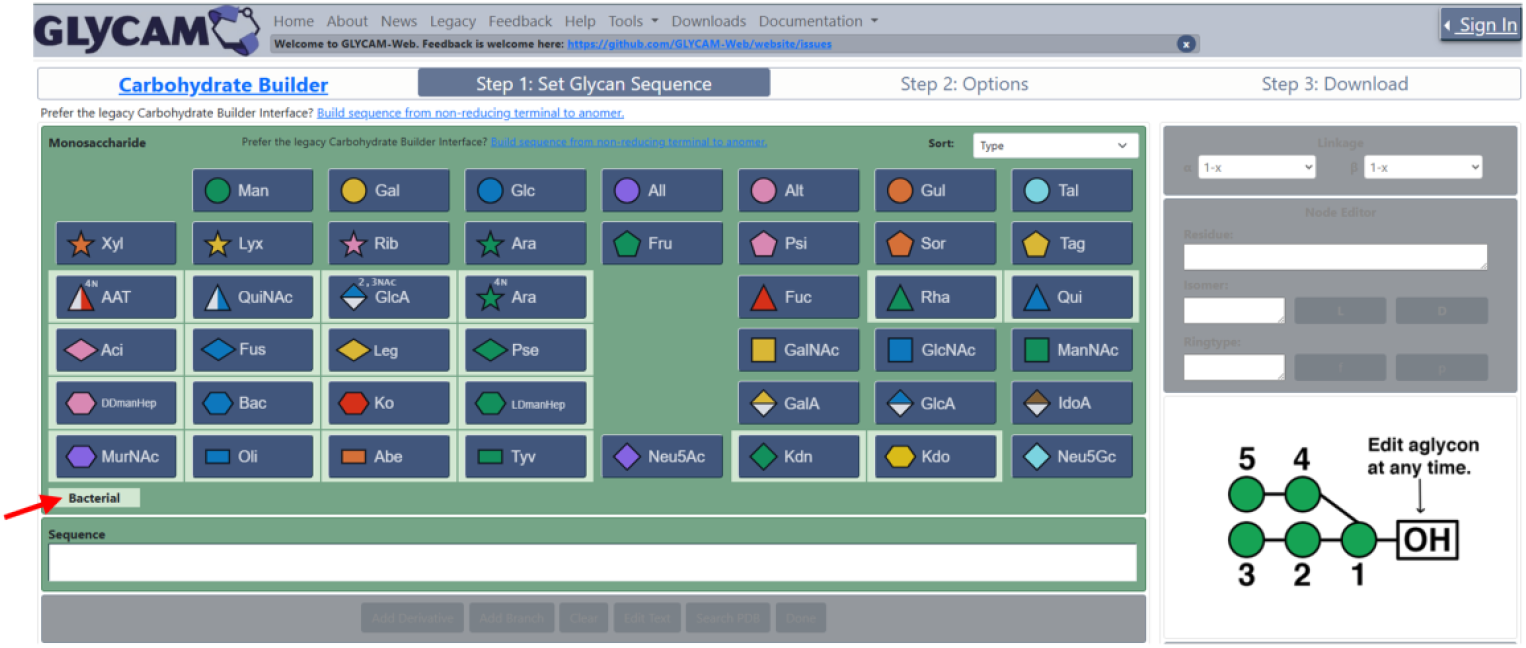
Illustration of the new point-and-click Glycam Bacterial Carbohydrate Builder interface. A light-green background (see red arrow) is used to visually distinguish the bacterial monosaccharide buttons from the rest of the library.

## RESULTS AND DISCUSSION

### GLYCAM Bacterial Carbohydrate Builder: scope and library composition

The newly developed GLYCAM Bacterial Carbohydrate Builder^1, 16^ empowers researchers to create 3D models of various bacterial glycans, addressing the unique challenges posed by their diversity and complexity. The initial set of parametrized sugars in GLYCAM Bacterial Carbohydrate Builder includes 18 selected bacterial monosaccharides, found as key components of bacterial glycoconjugates (Table 1, Figure 3), such as Kdo and Ko (abbreviated as K3O), heptoses, the uncommon aminosugar AAT (2-acetamido-4-amino-2,4,6-trideoxy-D-galactopyranose), Gal*f*, QuiNAc and Ara4N, mainly present in LPSs, as well as sialic acid-like sugars, including legionaminic, pseudaminic, acinetaminic and fusaminic acids, found in EPSs, CPSs and LPSs of several pathogens, and MurNAc, composing the saccharidic backbone of bacterial peptidoglycan.

**Table 1.**
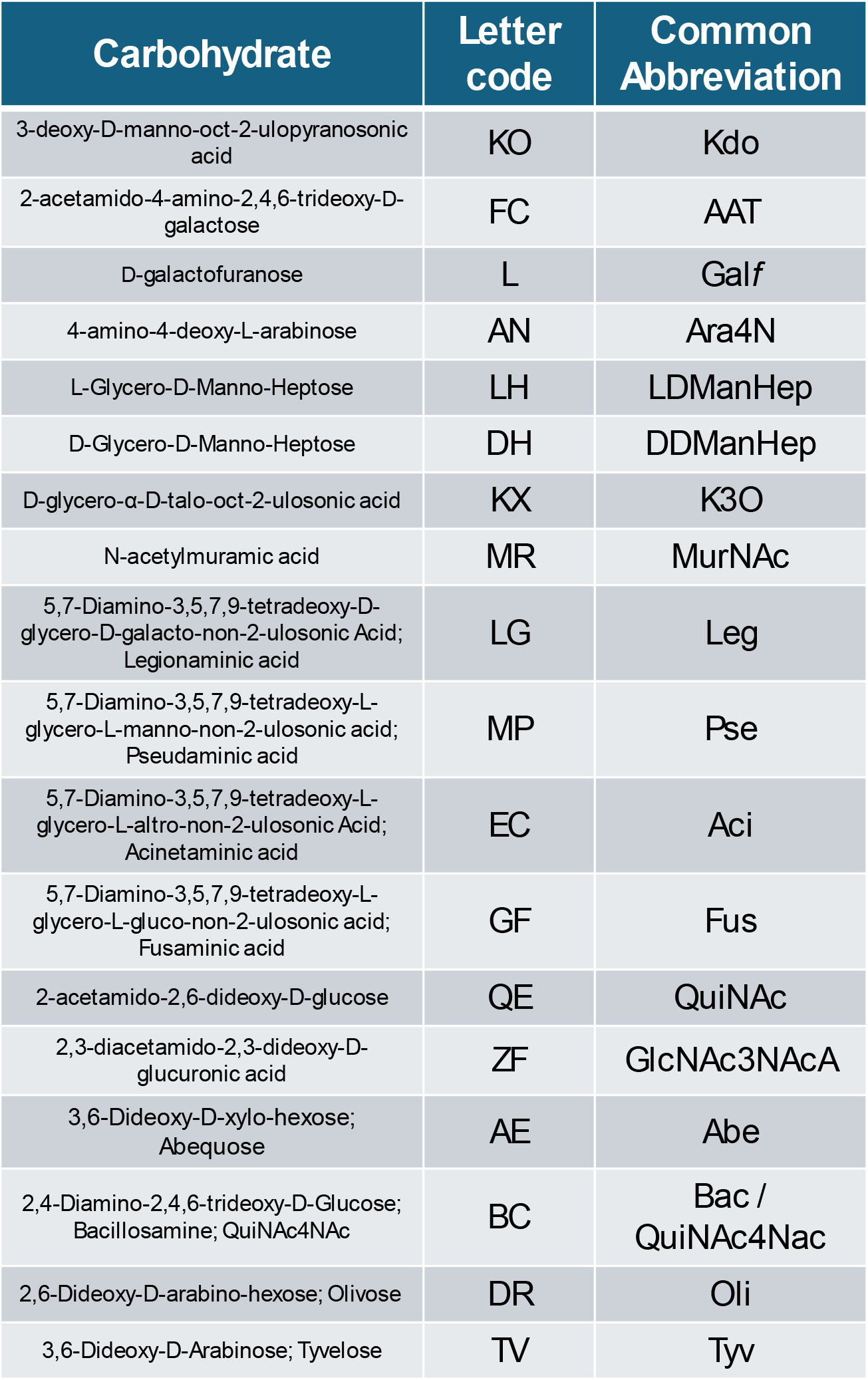
List of the 18 bacterial monosaccharides available on GLYCAM Bacterial Carbohydrate Builder. The full name, common abbreviations and corresponding letter codes are also reported.

**Figure 3.**
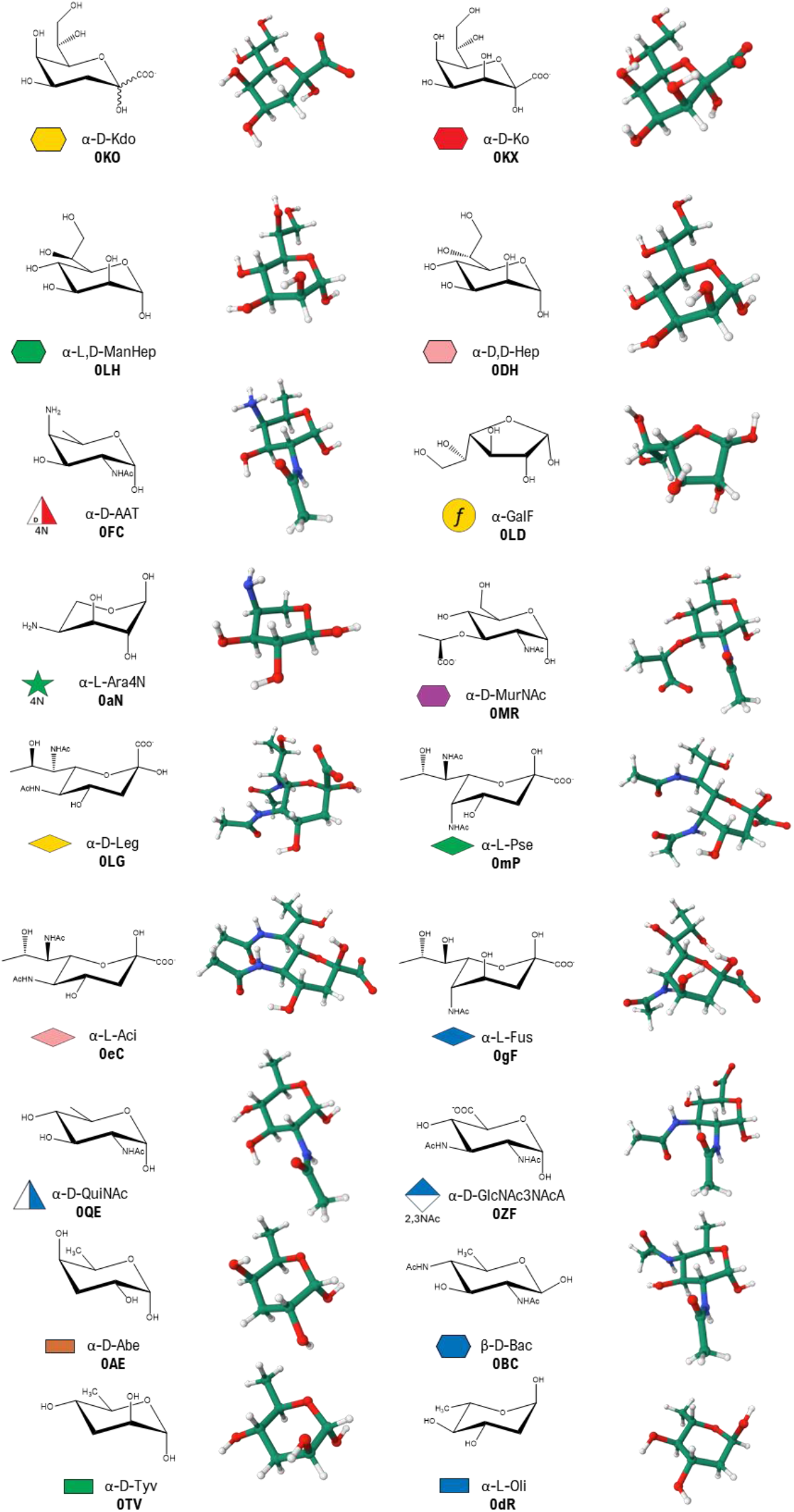
Chemical representation of the 18 bacterial monosaccharides available of GLYCAM Bacterial Carbohydrate Builder. The corresponding abbreviation and symbol in the SNFG (Symbol Nomenclature for Glycans) format are reported for each sugar.^17^

### Integration within GLYCAM-Web and structure generation workflow

GLYCAM Bacterial Carbohydrate Builder will provide users the possibility to mix bacterial and eukaryotic carbohydrates, offering flexibility for creating uncommon or novel glycans. This capability is crucial for researchers working with peculiar bacterial glycans. GLYCAM Bacterial Carbohydrate Builder allows users to select terminal residues and aglycons to complete their glycan structures,^18^ according to experimentally observed motifs, enhancing customization options. Once the glycan sequence is defined, the builder automatically performs energy minimisation of the structure using the GLYCAM06 force field,^19^ which includes bacterial glycan-specific parameters. The resulting models can be downloaded in PDB format and as input files suitable for molecular dynamics simulations using AMBER.^20^

In addition to sequence construction, chemical modifications commonly observed in bacterial monosaccharides can be introduced directly at the residue level. Supported derivatives include O-acetylation, methylation, sulfation, and deoxygenation, which can be applied to specific hydroxyl positions to reproduce experimentally characterised motifs found in bacterial glycoconjugates. These modifications are handled explicitly during model construction, with the “Add Derivative” button, which will allow to select the sugar, the position and the type of derivatization to add, ensuring consistency with the underlying GLYCAM06 parametrisation strategy.

### Generation of representative bacterial glycan models

As an alternative to the graphical carbohydrate builder, users may generate complete glycan structures using the “Build via Text” functionality (https://glycam.org/txt/, Figure 4). In this workflow, condensed sequences are provided as input, followed by sequence validation and optional refinement of glycosidic torsion angles prior to automated energy minimisation. This approach enables rapid generation of complex bacterial oligosaccharide structures without requiring manual assembly. Thus, to illustrate the utility of the builder, we provide representative examples of sequences, generated by using the presented tool, corresponding to the saccharidic region of peptidoglycan (PG) and lipopolysaccharides (LPS), as well as the repeating units of capsular polysaccharides (CPS), and exopolysaccharides (EPS). The corresponding condensed sequences are reported in Table 2 and can simply be copied and pasted into Step 1 (“Set Glycan Sequence”), and after optionally refining torsional angles in Step 2, the minimized coordinates can be downloaded in PDB format for further use in visualization or simulation workflows. The builder also allows users to add counter ions for charge neutrality and solvate the oligosaccharides with water, preparing them for realistic MD simulations.

**Table 2.**
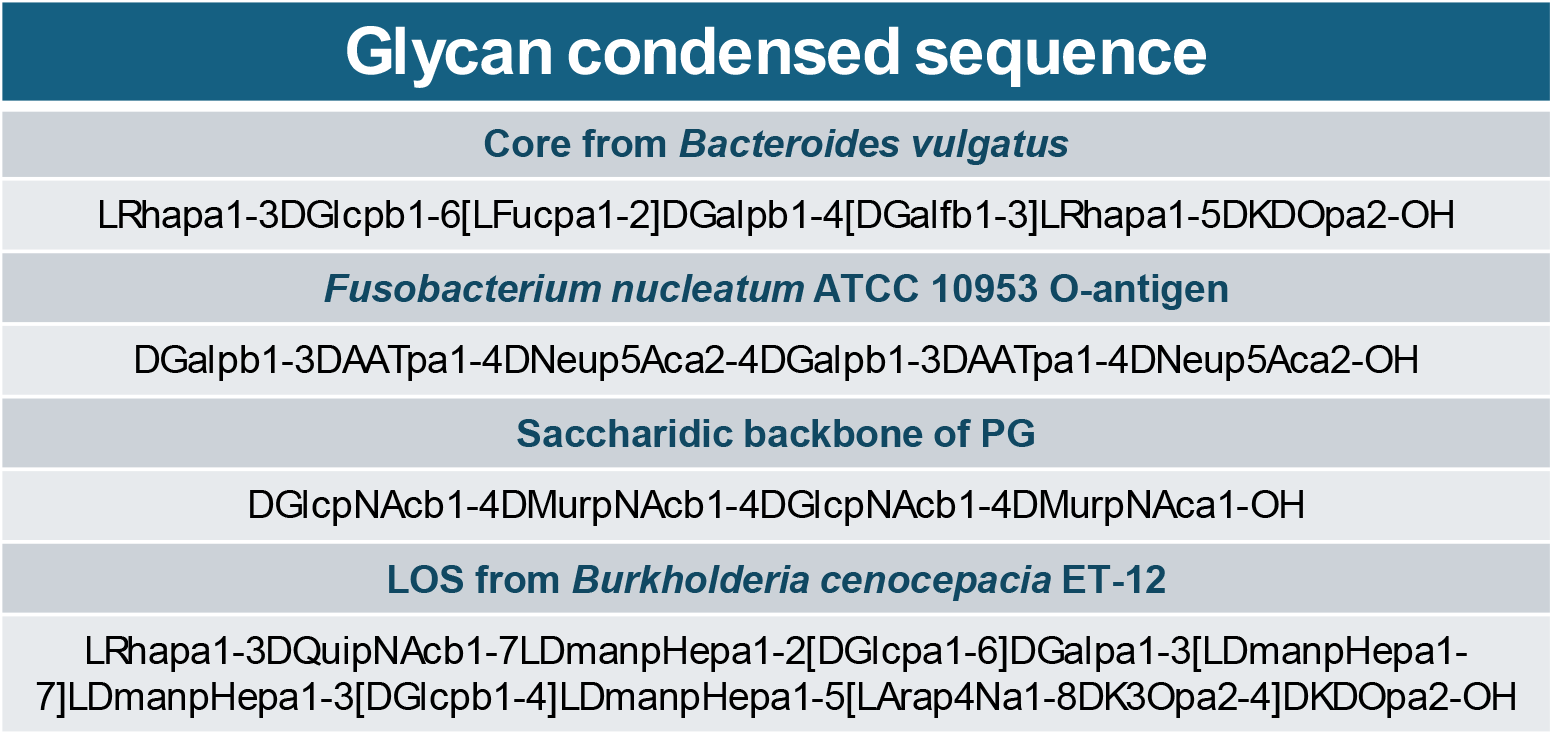
Representative bacterial glycan sequences for use with the “Build via Text” tool. Condensed sequences corresponding to selected bacterial glycans. Users can directly copy and paste the sequences into Step 1 of the “Build via Text” tool (https://glycam.org/txt/) to generate three-dimensional models (see Figure 4).

**Figure 4.**
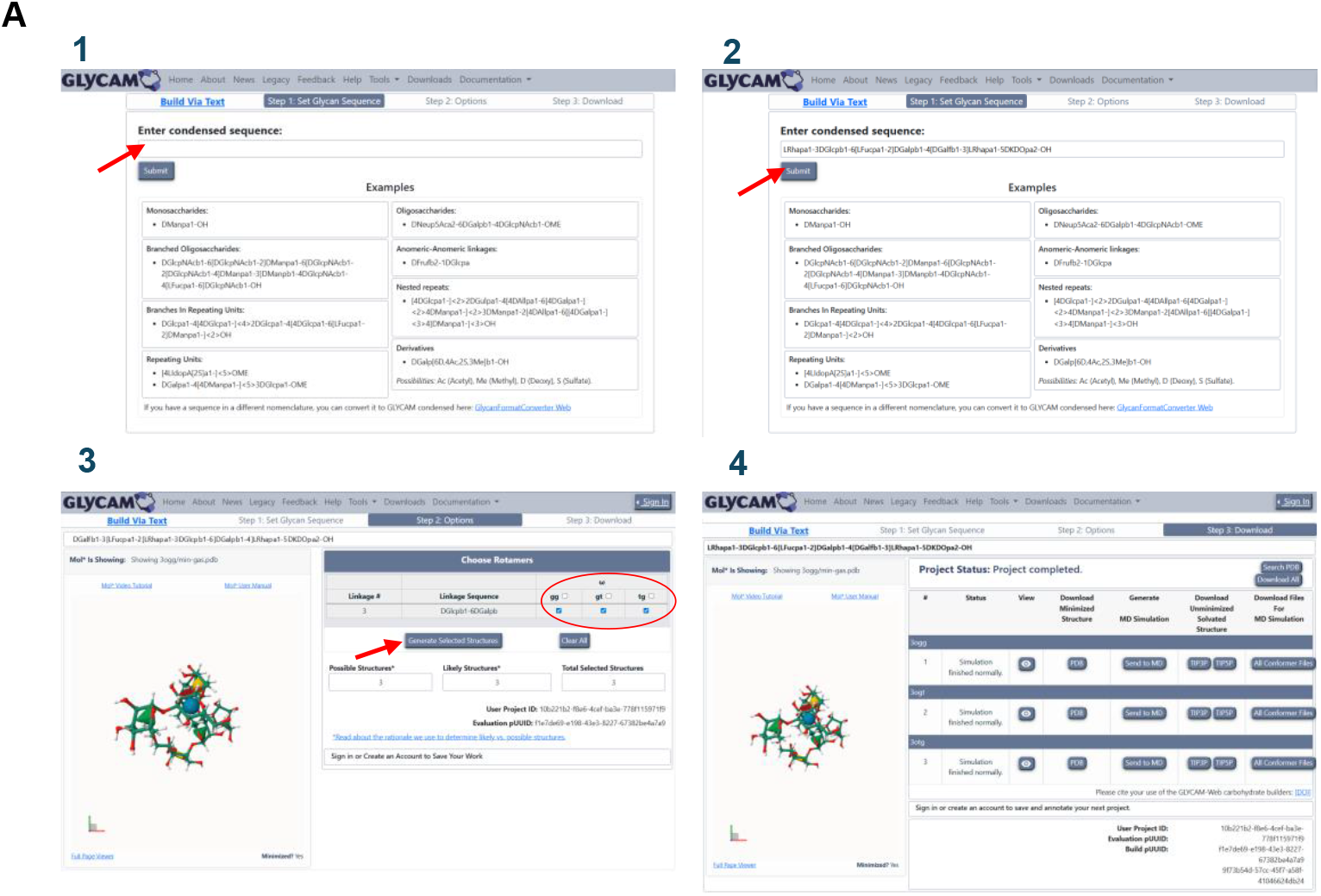
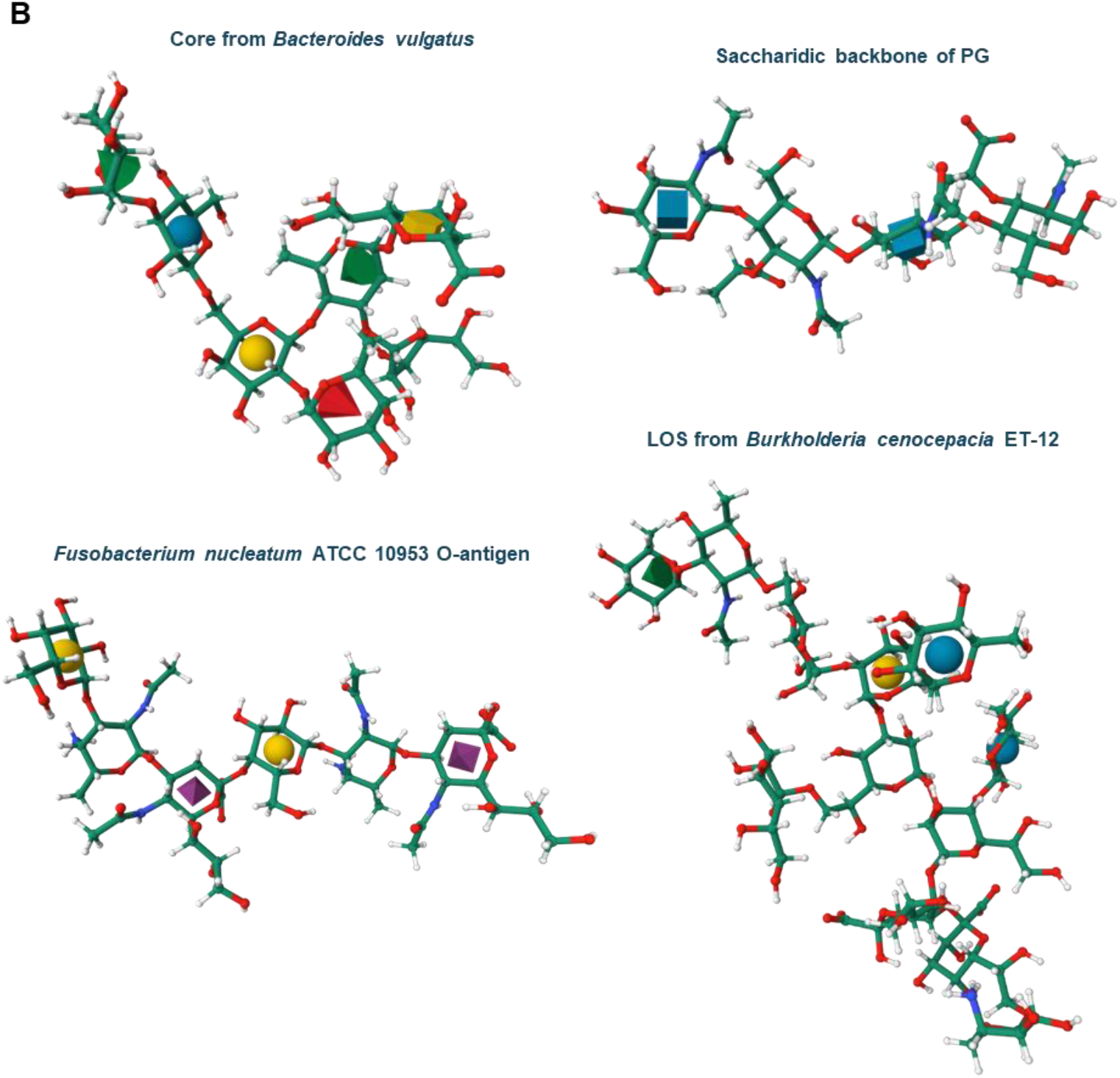
Representative workflow and examples generated with the GLYCAM Bacterial Carbohydrate Builder. **A)** Step-by-step procedure for constructing glycans using the “Build via Text” tool (https://glycam.org/txt/): (1) input of condensed sequence, (2) selection of glycan sequence, (3) optional refinement of torsional angles, and (4) energy minimization and download of the final structure. **B)** Representative 3D models obtained from sequences reported in Table 2, including the core from *Bacteroides vulgatus*,^8, 10^ the saccharidic backbone of peptidoglycan, a LOS from *Burkholderia cenocepacia* ET-12,^11^ and the O-antigen from *Fusobacterium nucleatum* ATCC 10953.^12^

### Validation through representative bacterial glycan case studies

To validate 3D models generated for bacterial glycans, it is essential to consider that carbohydrates are intrinsically flexible and are best described as conformational ensembles rather than single static structures.^18, 21^ Within this frame, MD simulations in aqueous environments are routinely employed to characterise several conformational states, allowing to identify the favoured torsions for each glycosidic linkage contained in the analysed 3D structure, and to assess their consistency with experimentally derived information (notably NMR data).^18, 22, 23^

To evaluate the performance and practical applicability of the GLYCAM Bacterial Carbohydrate Builder, we selected representative bacterial glycan systems spanning diverse glycoconjugate classes and structural complexities, including challenging oligosaccharide architectures. Accordingly, our validation strategy combines (i) reconstruction of representative bacterial glycan motifs using the public GLYCAM-Web implementation, (ii) automated GLYCAM06-based energy minimisation, and (iii) short MD-based checks of torsional preferences to benchmark against previously characterised systems and assess simulation readiness.

Representative bacterial glycan structures generated using the GLYCAM Bacterial Carbohydrate Builder were subjected to automated energy minimisation using the GLYCAM06 force field,^19^ following the same underlying framework employed in GLYCAM-Web^16^ for eukaryotic glycans. The resulting models were free of steric clashes and exhibited chemically reasonable geometries, enabling direct generation of AMBER-compatible MD input files without additional manual refinement. Importantly, all the bacterial glycan motifs considered here have been investigated previously by using experimental techniques (i.e. NMR spectroscopy) occasionally combined with computational studies. In the present work, we rebuild a couple of these motifs using the publicly available GLYCAM-Web implementation and benchmark the resulting structures against the conformational features established previously, thereby demonstrating that the public deployment reproduces the behaviour expected for chemically diverse bacterial glycans and provides simulation-ready starting conformations.

#### Bacteroides vulgatus mpk LPS core reconstruction and reproducibility assessment

As a first case study, we selected the lipopolysaccharide (LPS) of *Bacteroides vulgatus* mpk (BVMPK),^8, 10^ focusing on inner-core fragments that incorporate uncommon bacterial monosaccharide motifs not routinely available in standard eukaryotic glycan libraries. In BVMPK, these motifs include the Gram-negative signature residue 3-deoxy-D-manno-oct-2-ulopyranosonic acid (Kdo) and galactofuranose (Galf), providing a stringent test of bacterial carbohydrate modelling within the GLYCAM framework. Here, we rebuilt the BVMPK core heptasaccharide sequence (Table 2, Figure 4B) using the publicly available GLYCAM Bacterial Carbohydrate Builder, generating minimised PDB coordinates and AMBER-ready topology/coordinate files (PRMTOP/INPCRD) for direct use in MD simulations.

To assess simulation readiness and confirm reproducibility of conformational behaviour, the builder-generated heptasaccharide was solvated in explicit water and subjected to a standard minimization/heating/equilibration protocol followed by a 10 ns production MD run. After an initial relaxation phase, the glycan remained stable with no evidence of steric artefacts or linkage distortions, reaching an RMSD regime fluctuating around ∼3– 4 Å (Figure 5A). We next analysed glycosidic torsion-angle sampling (ϕ/ψ) for two diagnostic linkages involving the uncommon bacterial residues Kdo and Gal*f*. Both linkages populated well-defined torsional regions, as reflected by compact scatter distributions and their associated marginal ϕ and ψ histograms (Figure 5B–C). Importantly, the sampled torsional basins closely match those reported previously for the same BVMPK core motifs,^8, 10^ supporting the reproducibility of the public GLYCAM-Web deployment relative to earlier BVMPK conformational characterisation achieved from both experimental and computational studies.

**Figure 5.**
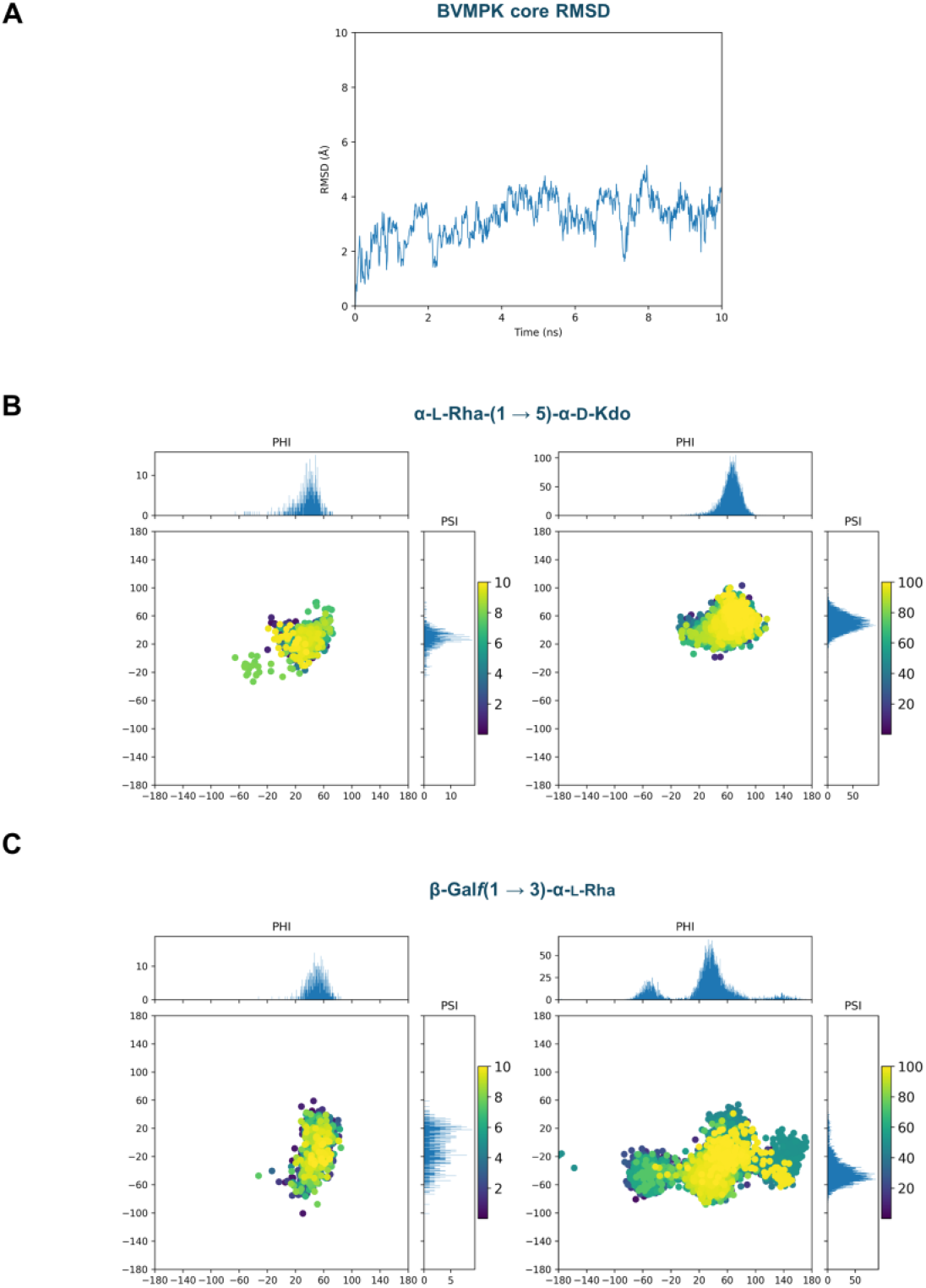
Public GLYCAM-Web reconstruction of the BVMPK LPS core reproduces previously reported conformation of the heptasaccharide, specifically including Kdo- and Galf-containing linkages, as derived from both experimental and computational studies.^8, 10^ **A)** RMSD of the BVMPK core heptasaccharide generated with the public GLYCAM Bacterial Carbohydrate Builder during a 10 ns explicit-solvent MD run, showing stable behaviour without structural artefacts. **B)** Glycosidic torsion-angle sampling (ϕ/ψ) for the α-L-Rha-(1→5)-α-D-Kdo linkage. The left panel shows the distribution obtained from the present work starting from the publicly generated model, whereas the right panel reports the corresponding distribution from the previously published BVMPK study. **C)** Glycosidic torsion-angle sampling (ϕ/ψ) for the β-Galf-(1→3)-α-L-Rha linkage. The left panel shows the distribution obtained from the present work starting from the publicly generated model, whereas the right panel reports the corresponding distribution from the previously published study. In panels B and C, marginal histograms show the ϕ and ψ projections; scatter points are coloured by simulation time (ns), illustrating consistent sampling patterns between the public GLYCAM-Web workflow and the reference trajectories.

#### Fusobacterium nucleatum Fn10953 O-antigen reconstruction and conformational reproducibility assessment

To complete the validation set, we examined the O-antigen of *Fusobacterium nucleatum ssp. polymorphum* ATCC 10953 (Fn10953).^12^ Whereas BVMPK probes Kdo- and Galf-containing core motifs, the Fn10953 O-antigen challenges the builder with a different chemical space: a sialylated repeating unit defined by a trisaccharide motif that incorporates the uncommon bacterial residue AAT (2-acetamido-4-amino-2,4,6-trideoxy-D-galactopyranose; also referred to as FucpNAc4N), linked to Neu5Ac and Gal (Table 2, Figure 4B).^12^ This system provides a stringent and chemically orthogonal benchmark because it combines a bacterial-specific amino sugar with Neu5Ac-containing linkages, offering a complementary test to the BVMPK core motifs. In the published Fn10953 study, oligomers containing increasing numbers of repeating units were characterised, and conformational analyses for the free-state 2-Mer identified distinct conformational families that can be used as a reference for modelling reproducibility.

As above, the corresponding oligosaccharide sequence was rebuilt using the public GLYCAM Bacterial Carbohydrate Builder, and the minimised output was carried forward into a standard explicit-solvent protocol followed by a short production MD run (10 ns). The glycan remained stable over the trajectory, with the RMSD rapidly equilibrating and fluctuating within a narrow range (Figure 6A), supporting the simulation readiness of the builder-derived starting conformation.

**Figure 6.**
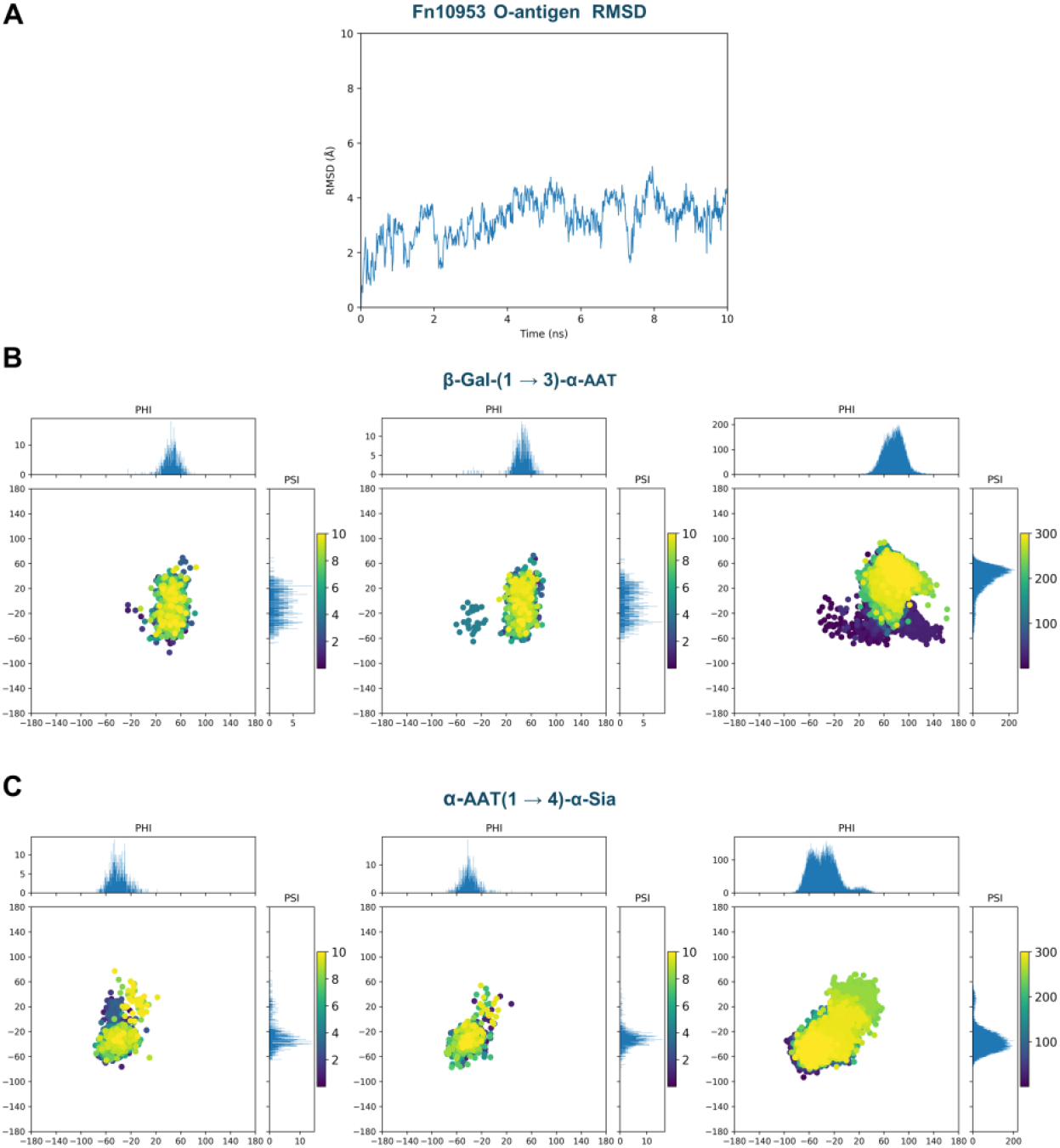
Public GLYCAM-Web reconstruction of the Fn10953 O-antigen reproduces previously reported conformation for AAT-containing linkages, as derived from both experimental and computational studies.^12^ A) RMSD of the Fn10953 O-antigen fragment generated with the public GLYCAM Bacterial Carbohydrate Builder during a 10 ns explicit-solvent MD run, indicating stable behaviour without evidence of artefacts. B) Glycosidic torsion-angle sampling (ϕ/ψ) for the β-Gal- (1→3)-α-AAT linkage. The left and centre panels report the two occurrences of this linkage within the simulated hexasaccharide (present work, starting from the publicly generated model), while the right panel reports the corresponding distribution from the previously published study. C) Glycosidic torsion-angle sampling (ϕ/ψ) for the α-AAT-(1→4)-α-Sia linkage. The left and centre panels report the two occurrences of this linkage within the simulated hexasaccharide (present work), while the right panel reports the corresponding distribution from the previously published study. In panels B and C, marginal histograms show the ϕ and ψ projections; scatter points are coloured by simulation time (ns), highlighting consistent sampling patterns between the public GLYCAM-Web workflow and the reference trajectories.

To assess conformational reproducibility relative to the previously established behaviour for this system, we examined the glycosidic torsion-angle sampling (ϕ/ψ) of the two linkages that define the Fn10953 repeating unit and involve the bacterial-specific residue AAT and the Neu5Ac-containing motif. For both linkages, the torsional distributions obtained from trajectories initiated from the public builder output reproduced the basins reported previously, with no evidence of sampling of sterically disfavoured regions (Figure 6B–C).

Together, these results indicate that the publicly deployed builder recapitulates the conformational preferences previously observed for the Fn10953 O-antigen motif while yielding directly simulation-ready models.

Overall, these case studies were chosen as representative stress tests rather than an exhaustive validation of the entire bacterial monosaccharide set. BVMPK and Fn10953 probe chemically and topologically demanding motifs (including Kdo, Galf, and AAT-containing linkages) for which prior conformational characterisation is available, enabling a direct reproducibility benchmark of the public GLYCAM-Web implementation. Beyond these in-depth benchmarks, all newly introduced bacterial residues in the current release undergo routine internal consistency checks within the GLYCAM06 framework (template integrity, stereochemistry, charge/type assignment, overlap resolution, and successful generation of AMBER-ready inputs), ensuring that the web-generated models are structurally sound and simulation-prepared.

### Limitations and future perspectives

The modelling of bacterial glycans is intrinsically challenged by the extraordinary diversity of bacterial monosaccharides and their chemical modifications. In contrast to eukaryotic systems, each bacterial sugar requires individual parametrisation to ensure compatibility with the GLYCAM06 force field. As a consequence, the current implementation of the GLYCAM Bacterial Carbohydrate Builder includes a curated subset of bacterial monosaccharides, selected based on their prevalence in well-characterised bacterial glycoconjugates. This initial library is intended as a starting point and will be progressively expanded, including new peculiar sugars as well as derivatizations, as additional parameters become available. Users interested in the parametrisation and inclusion of specific bacterial monosaccharides not currently supported are encouraged to contact the authors to discuss potential implementation within the GLYCAM framework. General limitations and methodological considerations inherent to the GLYCAM-Web building framework have been discussed in detail previously.^1^

We actively seek user feedback to enhance the functionality and accuracy of the GLYCAM Bacterial Carbohydrate Builder, ensuring it remains an indispensable tool for researchers studying the structure and function of bacterial glycans (https://glycam.org/docs/aboutus/contact-us/index.html). Our future plans include adding the 3D SNFG symbols for these bacterial sugars to Mol*^24^ the visualization software used on GLYCAM-Web that displays the 3D SNFG symbols^25, 26^

## METHODS

The GLYCAM Bacterial Carbohydrate Builder operates within the GLYCAM-Web framework and follows an established sequence-to-structure workflow for carbohydrate modelling.^16^ Glycan structures are generated from user-defined sequences, provided either through the graphical builder interface or via condensed text input. These sequences are parsed and translated into three-dimensional glycan topologies by sequential assembly of predefined monosaccharide templates according to their glycosidic connectivity and default conformational preferences.

During structure construction, potential atomic overlaps arising from branching patterns or unusual connectivities are resolved through automated adjustment of glycosidic torsion angles within experimentally supported ranges. The resulting structures are subsequently subjected to energy minimisation using the GLYCAM06 force field.^19^ Initial coordinate and topology files are generated in AMBER format and processed using the tleap^27^ module of AmberTools^27^ to produce PRMTOP and INPCRD files. Energy minimisation is then performed under dielectric conditions appropriate for aqueous environments. Following minimisation, the structures are analysed and processed using cpptraj,^28^ and, when requested by the user, explicitly solvated systems are generated using standard AMBER workflows. Solvation is performed using commonly employed three- or five-site water models, such as TIP3P^29^ or TIP5P^30^, and the corresponding coordinate and topology files are made available for download for subsequent molecular dynamics simulations.

Common chemical modifications observed in bacterial glycans, including O-acetylation, methylation, sulfation, and deoxygenation at selected positions, are supported through explicit residue-level handling during model construction, ensuring consistency with the underlying GLYCAM06 parametrisation strategy. A detailed description of the GLYCAM-Web modelling framework, structure assembly procedures, default conformational handling and its underlying algorithms has been reported previously.^1^

## ACKNOWLEDGMENTS

GLYCAM-Web was directly supported by NIH R24GM136984 (Transitioning Glycam-Web to a self-sustaining carbohydrate modeling service), U01CA207824 (Tools to enable nonspecialists) and R01GM100058 (Continued Development and Maintenance of GLYCAMWeb). Support from European Research Council (ERC) under the European Union”s Horizon 2020 research and innovation program under grant agreement No 851356 (R.M.). R. M. acknowledges HORIZON-MSCA-2023-DN-01 - GAP 101168287.

## REFERENCES

1. Grant, O. C.; Wentworth, D.; Holmes, S. G.; Kandel, R.; Sehnal, D.; Wang, X.; Xiao, Y.; Sheppard, P.; Grelsson, T.; Coulter, A.; Miller, G.; Foley, B. L.; Woods, R. J., Generating 3D Models of Carbohydrates with GLYCAM-Web. bioRxiv 2025, 2025.05.08.652828.

2. Nieto-Fabregat, F.; Marchetti, R.; Perez, S., Bacterial Polysaccharides : Integrated use of Databases and Computational Tools for their Structural Investigations. Perez, S., Ed. GlycoPedia, 2021.

3. Marchetti, R.; Forgione, R. E.; Fabregat, F. N.; Di Carluccio, C.; Molinaro, A.; Silipo, A., Solving the structural puzzle of bacterial glycome. Curr. Opin. Struct. Biol. 2021, 68, 74–83.

4. Vollmer, W.; Blanot, D.; De Pedro, M. A., Peptidoglycan structure and architecture. FEMS microbiology reviews 2008, 32 (2), 149–167.

5. Di Lorenzo, F.; Duda, K. A.; Lanzetta, R.; Silipo, A.; De Castro, C.; Molinaro, A., A Journey from Structure to Function of Bacterial Lipopolysaccharides. Chem. Rev. 2022, 122 (20), 15767–15821.

6. Di Lorenzo, F.; De Castro, C.; Silipo, A.; Molinaro, A., Lipopolysaccharide structures of Gram-negative populations in the gut microbiota and effects on host interactions. FEMS Microbiol Rev 2019, 43 (3), 257–272.

7. Cescutti, P., Chapter 6 - Bacterial capsular polysaccharides and exopolysaccharides. In Microbial Glycobiology, Holst, O.; Brennan, P. J.; Itzstein, M. v.; Moran, A. P., Eds. Academic Press: San Diego, 2010; pp 93–108.

8. Nieto-Fabregat, F.; Zhu, Q.; Vivès, C.; Zhang, Y.; Marseglia, A.; Chiodo, F.; Thépaut, M.; Rai, D.; Kulkarni, S. S.; Di Lorenzo, F.; Molinaro, A.; Marchetti, R.; Fieschi, F.; Xiao, G.; Yu, B.; Silipo, A., Atomic-Level Dissection of DC-SIGN Recognition of Bacteroides vulgatus LPS Epitopes. JACS Au 2024, 4 (2), 697–712.

9. Di Lorenzo, F.; Pither, M. D.; Martufi, M.; Scarinci, I.; Guzmán-Caldentey, J.; Łakomiec, E.; Jachymek, W.; Bruijns, S. C. M.; Santamaría, S. M.; Frick, J.-S.; van Kooyk, Y.; Chiodo, F.; Silipo, A.; Bernardini, M. L.; Molinaro, A., Pairing Bacteroides vulgatus LPS Structure with Its Immunomodulatory Effects on Human Cellular Models. ACS Cent. Sci. 2020, 6 (9), 1602–1616.

10. Rai, D.; Nieto-Fabregat, F.; Dikshit, R.; Thépaut, M.; Fieschi, F.; Silipo, A.; Kulkarni, S. S., Toward Glycomimetic Immunomodulators: A Synthetic Dissection of Phocaeicola vulgatus Core Oligosaccharides and Their Recognition by DC-SIGN. ACS Omega 2025, 10 (43), 51985–52000.

11. Silipo, A.; Molinaro, A.; Ieranò, T.; De Soyza, A.; Sturiale, L.; Garozzo, D.; Aldridge, C.; Corris, P. A.; Khan, C. M.; Lanzetta, R.; Parrilli, M., The complete structure and pro-inflammatory activity of the lipooligosaccharide of the highly epidemic and virulent gram-negative bacterium Burkholderia cenocepacia ET-12 (strain J2315). Chemistry 2007, 13 (12), 3501–11.

12. Di Carluccio, C.; Nieto-Fabregat, F.; Cerofolini, L.; Abreu, C.; Padilla-Cortés, L.; Gheorghita, G. R.; Masi, A. A.; Buono, L.; Gumah Adam Ali, M.; Lamprinaki, D.; Molinaro, A.; Juge, N.; Smaldone, G.; Vaněk, O.; Fragai, M.; Marchetti, R.; Silipo, A., Fusobacterium nucleatum Lipopolysaccharides O-Antigen Defines a Novel Siglec-7 Binding Epitope. JACS Au 2025.

13. Garcia-Vello, P.; Di Lorenzo, F.; Zucchetta, D.; Zamyatina, A.; De Castro, C.; Molinaro, A., Lipopolysaccharide lipid A: A promising molecule for new immunity-based therapies and antibiotics. Pharmacol Ther 2022, 230, 107970.

14. Vinogradov, E.; St. Michael, F.; Homma, K.; Sharma, A.; Cox, A. D., Structure of the LPS O-chain from Fusobacterium nucleatum strain 10953, containing sialic acid. Carbohydr. Res. 2017, 440-441, 38–42.

15. De Castro, C.; Holst, O.; Lanzetta, R.; Parrilli, M.; Molinaro, A., Bacterial lipopolysaccharides in plant and mammalian innate immunity. Protein and peptide letters 2012, 19 (10), 1040–4.

16. Group, W. GLYCAM Web. http://glycam.org.

17. Neelamegham, S.; Aoki-Kinoshita, K.; Bolton, E.; Frank, M.; Lisacek, F.; Lütteke, T.; O”Boyle, N.; Packer, N. H.; Stanley, P.; Toukach, P.; Varki, A.; Woods, R. J.; The, S. D. G., Updates to the Symbol Nomenclature for Glycans guidelines. Glycobiology 2019, 29 (9), 620–624.

18. Woods, R. J., Predicting the Structures of Glycans, Glycoproteins, and Their Complexes. Chemical Reviews 2018, 118 (17), 8005–8024.

19. Kirschner, K. N.; Yongye, A. B.; Tschampel, S. M.; González-Outeiriño, J.; Daniels, C. R.; Foley, B. L.; Woods, R. J., GLYCAM06: a generalizable biomolecular force field. Carbohydrates. J Comput Chem 2008, 29 (4), 622–55.

20. Case, D. A.; Aktulga, H. M.; Belfon, K.; Ben-Shalom, I. Y.; Berryman, J. T.; Brozell, S. R.; Carvahol, F. S.; Cerutti, D. S.; Cheatham, T. E.; Cisneros, G. A.; Cruzeiro, V. W. D.; Darden, T. A.; Forouzesh, N.; Ghazimirsaeed, M.; Giambaşu, G.; Giese, T.; Gilson, M. K.; H. Gohlke; A.W. Goetz; Harris, J.; Huang, Z.; Izadi, S.; Izmailov, S. A.; Kasavajhala, K.; Kaymak, M. C.; Kolossvary, I.; Kovalenko, A.; Kurtzman, T.; Lee, T. S.; Li, P.; Li, Z.; Lin, C.; Liu, J.; Luchko, T.; Luo, R.; Machado, M.; Manathunga, M.; Merz, K. M.; Miao, Y.; Mikhailovskii, O.; Monard, G.; Nguyen, H.; O”Hearn, K. A.; Onufriev, A.; Pan, F.; Pantano, S.; Rahnamoun, A.; Roe, D. R.; Roitberg, A.; Sagui, C.; Schott-Verdugo, S.; Shajan, A.; Shen, J.; Simmerling, C. L.; Skrynnikov, N. R.; Smith, J.; Swails, J.; Walker, R. C.; Wang, J.; Wang, J.; Wu, X.; Wu, Y.; Xiong, Y.; Xue, Y.; York, D. M.; Zhao, C.; Zhu, Q.; Kollman, P. A. AMBER 2018, University of California, San Francisco, 2025.

21. Fadda, E.; Woods, R. J., Molecular simulations of carbohydrates and protein– carbohydrate interactions: motivation, issues and prospects. Drug Discovery Today 2010, 15 (15), 596–609.

22. Zhang, W.; Turney, T.; Meredith, R.; Pan, Q.; Sernau, L.; Wang, X.; Hu, X.; Woods, R. J.; Carmichael, I.; Serianni, A. S., Conformational Populations of β-(1→4) O-Glycosidic Linkages Using Redundant NMR J-Couplings and Circular Statistics. The Journal of Physical Chemistry B 2017, 121 (14), 3042–3058.

23. Meredith, R. J.; Carmichael, I.; Woods, R. J.; Serianni, A. S., MA”AT Analysis: Probability Distributions of Molecular Torsion Angles in Solution from NMR Spectroscopy. Accounts of Chemical Research 2023, 56 (17), 2313–2328.

24. Sehnal, D.; Bittrich, S.; Deshpande, M.; Svobodová, R.; Berka, K.; Bazgier, V.; Velankar, S.; Burley, S. K.; Koča, J.; Rose, A. S., Mol* Viewer: modern web app for 3D visualization and analysis of large biomolecular structures. Nucleic Acids Research 2021, 49 (W1), W431–W437.

25. Sehnal, D.; Grant, O. C., Rapidly Display Glycan Symbols in 3D Structures: 3D-SNFG in LiteMol. Journal of Proteome Research 2019, 18 (2), 770–774.

26. Varki, A.; Cummings, R. D.; Aebi, M.; Packer, N. H.; Seeberger, P. H.; Esko, J. D.; Stanley, P.; Hart, G.; Darvill, A.; Kinoshita, T.; Prestegard, J. J.; Schnaar, R. L.; Freeze, H. H.; Marth, J. D.; Bertozzi, C. R.; Etzler, M. E.; Frank, M.; Vliegenthart, J.F. G.; Lütteke, T.; Perez, S.; Bolton, E.; Rudd, P.; Paulson, J.; Kanehisa, M.; Toukach, P.; Aoki-Kinoshita, K. F.; Dell, A.; Narimatsu, H.; York, W.; Taniguchi, N.; Kornfeld, S., Symbol Nomenclature for Graphical Representations of Glycans. Glycobiology 2015, 25 (12), 1323–1324.

27. Case, D. A.; Aktulga, H. M.; Belfon, K.; Cerutti, D. S.; Cisneros, G. A.; Cruzeiro, V. W. D.; Forouzesh, N.; Giese, T. J.; Götz, A. W.; Gohlke, H.; Izadi, S.; Kasavajhala, K.; Kaymak, M. C.; King, E.; Kurtzman, T.; Lee, T.-S.; Li, P.; Liu, J.; Luchko, T.; Luo, R.; Manathunga, M.; Machado, M. R.; Nguyen, H. M.; O”Hearn, K. A.; Onufriev, A. V.; Pan, F.; Pantano, S.; Qi, R.; Rahnamoun, A.; Risheh, A.; Schott-Verdugo, S.; Shajan, A.; Swails, J.; Wang, J.; Wei, H.; Wu, X.; Wu, Y.; Zhang, S.; Zhao, S.; Zhu, Q.; Cheatham, T. E., III; Roe, D.R.; Roitberg, A.; Simmerling, C.; York, D. M.; Nagan, M.C.; Merz, K. M., Jr., AmberTools. Journal of Chemical Information and Modeling 2023, 63 (20), 6183–6191.

28. Roe, D. R.; Cheatham III, T. E., PTRAJ and CPPTRAJ: software for processing and analysis of molecular dynamics trajectory data. J. Chem. Theory Comput. 2013, 9 (7), 3084–3095.

29. Jorgensen, W. L.; Chandrasekhar, J.; Madura, J. D.; Impey, R. W.; Klein, M. L., Comparison of simple potential functions for simulating liquid water. The Journal of Chemical Physics 1983, 79 (2), 926–935.

30. Mahoney, M. W.; Jorgensen, W. L., A five-site model for liquid water and the reproduction of the density anomaly by rigid, nonpolarizable potential functions. The Journal of Chemical Physics 2000, 112 (20), 8910–8922.

